# Epilepsy-associated increase in gonadotropin-releasing hormone neuron firing in diestrous female mice is independent of chronic seizure burden severity

**DOI:** 10.1101/2022.02.13.480255

**Authors:** Jiang Li, Catherine A. Christian-Hinman

## Abstract

Reproductive endocrine disorders are common comorbidities of temporal lobe epilepsy (TLE). Our previous studies using the intrahippocampal kainic acid (IHKA) mouse model of TLE demonstrated that many females show prolonged estrous cycles and hypothalamic gonadotropin-releasing hormone (GnRH) neurons exhibit elevated firing during diestrus. However, it is unknown whether the degree of change in GnRH neuron activity is dependent on epilepsy severity. Here, we used 24/7 *in vivo* EEG and *in vitro* electrophysiological recordings in acute brain slices to assess GnRH neuron firing in relation to chronic seizure burden in diestrous female mice at two months after IHKA injection. We found that percentage of time in seizure activity in the 24 hours prior to slice preparation is an accurate proxy of overall seizure burden. Firing rates of GnRH neurons from EEG-recorded IHKA mice were increased in comparison to controls, but no relationships were found between GnRH neuron firing and seizure burden measured *in vivo*. The independence of GnRH neuron firing rate in relation to seizure burden was unaffected by GnRH neuron soma location or estrous cycle length. Furthermore, GnRH neuron firing rates were not yet different from control values when measured 1 month after injection when epileptogenesis is already complete in IHKA mice. This result suggests a potential window for intervention to prevent epilepsy-associated GnRH neuron dysfunction. These findings indicate that the severity of epilepsy and the degree of downstream disruption to GnRH neuron activity are independent, suggesting that susceptibility to reproductive endocrine comorbidities is driven by other risk factors.

## 1. Introduction

Reproductive endocrine disorders are common comorbidities of epilepsy, particularly temporal lobe epilepsy (TLE). In women, these comorbidities include polycystic ovary syndrome, hypothalamic amenorrhea, and premature menopause (Bauer, 2001; Herzog, 2008; Kalinin and Zheleznova, 2007; Klein et al., 2001; Nappi et al., 1994; Pennell, 2009; Penovich, 2000). Although some reproductive endocrine comorbidities arise as side effects of antiseizure drugs (Bauer et al., 2002; Isojärvi et al., 2001, 1993; Stoffel-Wagner et al., 1998), a large degree of reproductive endocrine dysfunction appears to arise from seizure activity itself (Bauer et al., 2000; Herzog, 2008, 1993; Herzog et al., 1986; Kalinin and Zheleznova, 2007). Increased seizure frequency has been linked to earlier age of menopause in women with epilepsy (Harden et al., 2003), although other studies have not found this association (Klein et al., 2001). In addition, a longer window of seizure exacerbation during the luteal phase in anovulatory menstrual cycles (Herzog et al., 2011, 1997; Kim et al., 2010) indicates a potential association between the severity of epilepsy and downstream reproductive endocrine dysfunction. The factors that initiate and determine the degree of comorbid dysfunction, however, remain unclear.

Clinical evidence for reproductive endocrine comorbidities in patients with TLE is recapitulated in preclinical studies examining various rodent models of TLE (Christian et al., 2020). Disruption of the female estrous cycle, the rodent correlate of the human menstrual cycle, has been documented following systemic pilocarpine-induced status epilepticus in rats and mice (Fawley et al., 2012; Scharfman et al., 2008), and amygdala kindling (Edwards et al., 1999) and intrahippocampal kainic acid (IHKA) injection in rats (Amado et al., 1987). Furthermore, we previously reported that in the IHKA mouse model of TLE, many female mice, but not all, show prolonged estrous cycles and disrupted hypothalamic control of reproduction (Li et al., 2018, 2017). Furthermore, we found that hypothalamic gonadotropin-releasing hormone (GnRH) neurons exhibit elevated firing rates during diestrus and diminished activity during estrus in IHKA female mice when measured two months after IHKA injection (Li et al., 2018). GnRH neurons are the final common output pathway from the brain in the neural control of reproduction through the hypothalamic-pituitary-gonadal (HPG) axis. Therefore, understanding the physiological parameters that shape the response of GnRH neuron activity to the challenge of epilepsy should provide mechanistic insight into the causes and development of epilepsy-associated reproductive endocrine comorbidities.

One factor that could shape development of certain comorbidities is epilepsy severity, particularly considering the high degree of heterogeneity in seizure properties and frequency among patients with TLE (Bartolomei et al., 2008; de Barros Lourenço et al., 2020; Maillard et al., 2004). It is unclear whether seizure burden and comorbid reproductive endocrine disorders are correlated clinically, but seizure burden is positively correlated with comorbid depression (Cramer et al., 2003) and cognitive impairments (Holmes, 2015). Understanding whether epilepsy severity influences the propensity to develop comorbid reproductive endocrine problems is vital to identify potential underlying mechanisms. Therefore, the objective of this study was to determine whether the severity of chronic epilepsy, as represented by electroencephalography (EEG)-recorded seizure burden, displays correlated relationships to GnRH neuron firing rate in the IHKA mouse model of epilepsy.

The IHKA mouse model is advantageous for this investigation because the mice naturally develop a strong heterogeneity in seizure burden from the same targeted injection (Bouilleret et al., 1999; Li et al., 2020; Rusina et al., 2021; Zeidler et al., 2018), recapitulating the variability in epilepsy severity in TLE. Moreover, female IHKA mice demonstrate differential estrous cycle phenotypes, with some developing long, disrupted cycles and others maintaining regular cycles (Li et al., 2018, 2017). Therefore, in the present studies we assessed GnRH neuron function in relation to the severity of epilepsy in IHKA female mice. Only females were used as the experimental goal was to determine if our previous sex-specific finding of increased GnRH neuron firing in diestrous IHKA females with long estrous cycles (Li et al., 2018) is correlated to the degree of EEG-recorded seizure burden. In addition, to begin to determine the time course of disruption of GnRH neuron function in relation to the development of epilepsy in this model, we also recorded GnRH neuron firing at one month after injection, when spontaneous seizures are typically already present in IHKA mice (Bouilleret et al., 1999; Gröticke et al., 2008; Lee et al., 2012; Riban et al., 2002).

## 2. Materials and Methods

### 2.1. Animals

All animal procedures were approved by the Institutional Animal Care and Use Committee of the University of Illinois Urbana-Champaign and conducted in accordance with the ARRIVE guidelines. EEG-recorded animals used in this study were included in a previous report; details of animal housing, transgenic mouse breeding, estrous cycle monitoring procedures, IHKA injection and EEG electrode implantation surgical procedures, and video-EEG recording methods can be found in that publication (Li et al., 2020). Briefly, GnRH-tdTomato mice were produced by a cross of GnRH-Cre (Yoon et al., 2005) and Ai9 (Madisen et al., 2010) strains. Daily estrous cycle monitoring by vaginal cytology (Pantier et al., 2019) started on postnatal day (P) 42. After at least 2 weeks of monitoring, females with regular estrous cycles underwent intrahippocampal injection of either KA (Tocris; 50 nl of 20 mM, n = 9 mice) or control sterile 0.9% saline (n = 6 mice) on diestrus, followed by bipolar EEG electrode implantation in the ipsilateral hippocampus approximately 2 weeks later. Both surgical procedures were performed under isoflurane anesthesia. Mice were recorded in 24/7 video-EEG using recorder software written in MATLAB (Armstrong et al., 2013; Zeidler et al., 2018) from 21-35 days after injection (1 month) and again from 49-70 days after injection (2 months). A separate cohort of 10 GnRH-tdTomato females underwent intrahippocampal injection of KA (n = 6) or saline (n = 4) on diestrus without subsequent EEG implantation or recording.

### 2.2. Brain slice preparation

On a day of diestrus approximately 2 months after intrahippocampal injection, mice were disconnected from EEG recording and coronal brain slices were prepared as previously described (Li et al., 2018) after euthanasia by decapitation between Zeitgeber time (ZT) 2-3 (relative to ZT 12 lights-off). Brain slice preparation for mice not subjected to EEG recording was done in the same manner at 1 month after injection. 300 µm-thick coronal brain slices were made in ice-cold oxygenated sucrose solution (in mM: 2.5 KCl, 1.25 NaH_2_PO_4_, 10 MgSO_4_, 0.5 CaCl_2_, 11 glucose, and 234 sucrose) using a Leica Biosystems VT1200S vibrating blade microtome, then transferred to oxygenated artificial cerebrospinal fluid (ACSF; in mM: 2.5 KCl, 10 glucose, 126 NaCl, 1.25 NaH_2_PO_4_, 1 MgSO_4_, 0.5 CaCl_2_, and 26 NaHCO_3_; osmolarity 298 mOsm) for 30 min at 32° C before being transferred to room temperature (∼22-24° C) for at least 30 min prior to recording.

### 2.3. Targeted extracellular recordings

Action currents in tdTomato-identified GnRH neurons were recorded as described previously (Li et al., 2018). Briefly, individual slices were placed in a submerged recording chamber on the stage of an upright BX51WI microscope (Olympus). Oxygenated bath ACSF was superfused through the slice chamber at a flow rate of 2.5 ml/min and warmed to 30-32° C using an inline heater (Warner Instruments). Borosilicate glass recording pipettes (∼2 MΩ tip resistance) were prepared using an electrode puller (P-1000, Sutter Instruments) and filled with filtered ACSF solution with 10 mM HEPES buffer added. Each GnRH neuron was recorded for at least 20 min in voltage-clamp mode with holding potential set at 0 mV and signals Bessel-filtered at 12 kHz using a MultiClamp 700B amplifier, Digidata 1550 digitizer, and Clampex 10 software (Molecular Devices). Action current detection was done using Clampfit 10.6 software. GnRH neurons in the medial septum (MS) and preoptic area (POA) were identified and targeted for recording based on previous findings indicating that increased GnRH neuron activity in diestrous IHKA mice was limited to neurons with somata in these regions (Li et al., 2018). To evenly sample GnRH neuron firing in the MS and POA, at least one neuron per mouse was recorded at each location. Mean firing rate for each neuron (in Hz) was calculated as the number of action currents detected divided by the recording time in seconds. If no spontaneous action currents were detected, 15 mM KCl was added to the bath to induce firing and confirm successful recording. No more than 4 total GnRH neurons per mouse were included in analyses.

### 2.4. EEG analysis

Seizures were detected by a custom convolutional neural network machine learning model as previously described (Li et al., 2020), with seizure detection parameters set at minimum 6-second duration, 5 seconds inter-seizure interval, and seizure end delineated by reduction of spiking below 2 Hz. Only mice with EEG-confirmed epilepsy (defined as at least 2 spontaneous seizures) were included in further analyses. For mice with <1% of time in seizure activity detected in EEG analysis, seizures and other abnormal behaviors were confirmed in video recordings to identify and distinguish epileptiform events from false-positive artifacts.

### 2.5. Statistics

Statistical analyses were performed using OriginPro (OriginLab, Northampton, MA). Based on data normality as tested using Shapiro-Wilk tests, group comparisons between saline and IHKA groups were performed using nonparametric Mann-Whitney tests. Correlations were tested using Pearson or Spearman correlations as specified. p<0.05 was considered significant.

## 3. Results

### 3.1. Percentage of time in seizures in last 24 hours of recording accurately approximates overall seizure burden

There are multiple parameters of chronic seizure activity, including the percentage of time in seizure activity, seizure frequency (defined here as number of seizures per hour), and seizure duration, that can be used to quantify chronic seizure burden and epilepsy severity. To determine whether percentage of time in seizure activity is a robust representation of overall seizure burden, we assessed the relationships between percentage of time in seizure activity to number of seizures per hour and average seizure duration on all recorded days of diestrus. Strong positive correlations were found between percentage of time spent in seizures and both seizure frequency (r^2^ = 0.85, p = 0.00028, Pearson correlation) and average seizure duration (r^2^= 0.63, p = 0.0065, Pearson correlation; **Fig. 1**), indicating that the percentage of time in seizure activity is a suitable proxy for overall chronic seizure burden in these mice.

**Fig. 1.**
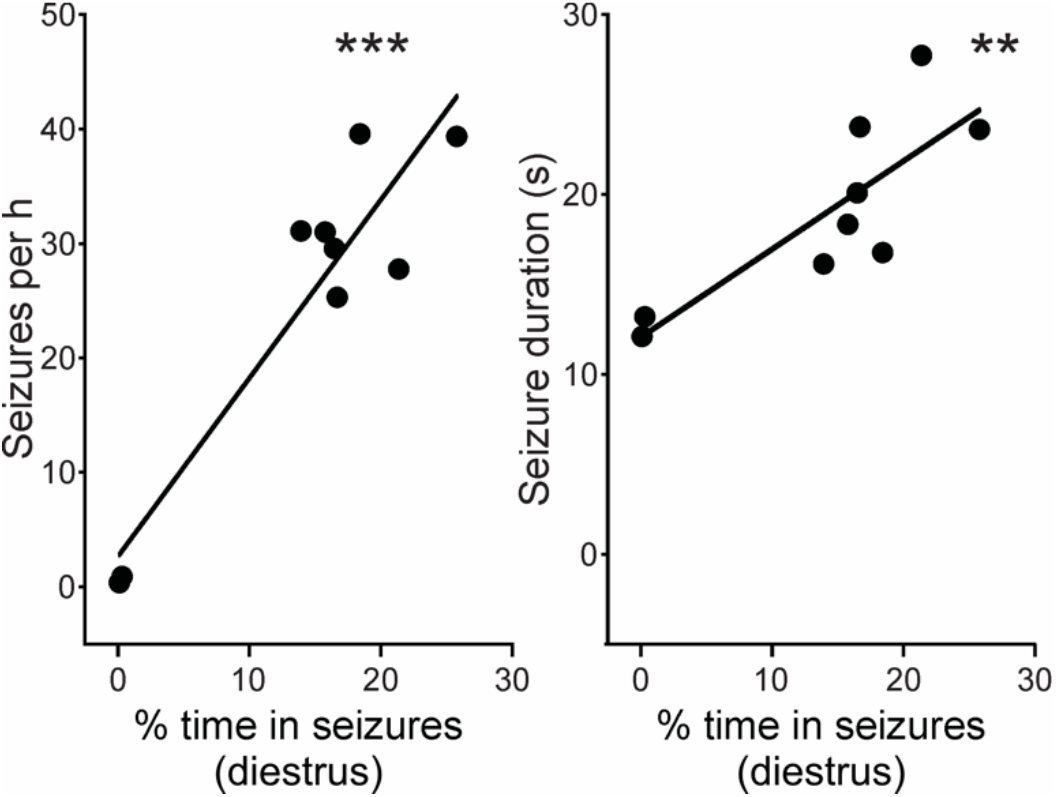
Percentage of time spent in seizure activity is strongly correlated to seizure frequency (left) and duration (right) on diestrus. Black circles indicate individual IHKA mice (n=9). Solid lines indicate the linear regression of best fit. **p<0.01, ***p<0.001 Pearson correlations.

To identify the window of seizure recording time most appropriate for correlations to subsequent *in vitro* analyses, we assessed the stability of chronic seizure burden in IHKA female mice at approximately 2 months after IHKA injection by performing a retrospective analysis of percentage of time in seizure activity recorded in the 9 hours (ZT 17-2; 0000-0900 h), 24 hours (ZT 2 – ZT 2; 0900 h – 0900 h), and 7 days before the mice were disconnected from the EEG recording system at ZT 2 on a day of diestrus. Strong correlations in percentage of time spent in seizures were observed between each of the time windows (9 hours vs. 24 hours, r^2^ = 0.98, p = 9.04×10^−8^; 9 hours vs. 7 days, r^2^ = 0.99, p = 3.51×10^−8^; 24 hours vs. 7 days, r^2^ = 0.98, p = 1.62×10^−7^, Pearson correlations; **Fig. 2A-C**). These findings indicated that percentage of time in seizure activity was stable across several time sampling windows ranging from 9 hours to 7 days. Therefore, for further analyses as described below, we used the values obtained for the 24-hour period prior to the end of EEG recording as a reasonable approximation of overall seizure burden in each mouse.

**Fig. 2.**
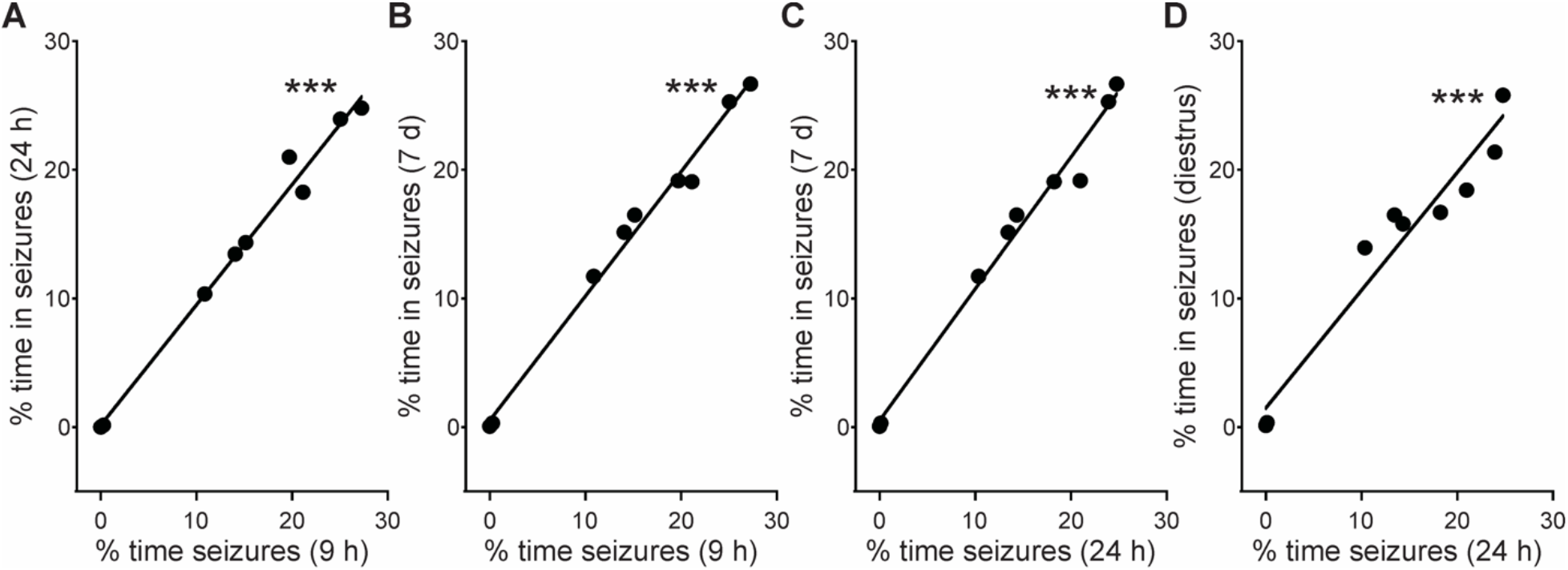
Strong correlations between values for percentage of time in seizure activity quantified for 9 hours, 24 hours, and 1 week prior to cessation of EEG recording on diestrus. A-C) Relationship of percentage of time in seizures in last 9 hours recorded as a function of values for last 24 hours (A), in last 9 hours as a function of values for last week (B), and for last 24 hours as a function of values for last week (C). D) Relationship of percentage of time in seizures in last 24 hours as a function of values for all days of diestrus collected. Black circles indicate individual IHKA mice (n=9). Solid lines indicate the linear regressions of best fit. ***p<0.001, Pearson correlations

Our previous analysis of EEG patterns, which included the mice examined here, indicated the presence of fluctuations in seizure burden with estrous cycle stage, with increased seizure burden on proestrus and estrus compared with diestrus (Li et al., 2020). The time windows examined in the analyses in Fig. 2A-C were all relative to a day of diestrus, and diestrus is typically preceded either by metestrus or another day of diestrus. Therefore, we analyzed the correspondence between the proximal seizure burden the mice experienced in the 24 hours prior to the end of EEG recording and the levels of seizure burden detected on all days of diestrus that were collected during EEG recordings from 49-70 days after injection. The percentage of time in seizure activity leading up to the last day of recording strongly correlated with the percentage of time in seizures calculated from all diestrus days (r^2^ = 0.93, p = 1.37×10^− 5^, Pearson correlation; **Fig. 2D**). This result further suggested that the last 24 hours of EEG recording provided a strong approximation of the seizure burden each mouse would have experienced in the period during which subsequent *in vitro* slice electrophysiology recordings were performed, as described below.

### 3.2. Lack of correlation between seizure burden and mean GnRH neuron firing rate in individual mice

Our previous report described increased firing activity of GnRH neurons from IHKA mice on diestrus (Li et al., 2018), but the mice in that study were not recorded in EEG. Here, we compared the firing rates of GnRH neurons in the MS and POA from saline-injected control or IHKA mice, all subjected to the same regimen of EEG tethering and recording. In this cohort, GnRH neurons from IHKA mice (n=27 cells total; MS n=15, POA n=12) exhibited increased firing rates compared to neurons from control mice (n=19 cells total; MS n=8, POA n=11; p = 0.0046, Mann-Whitney test) (**Fig. 3A-B**). These data thus replicate our previous finding of increased activity of MS and POA GnRH neurons in IHKA females (Li et al., 2018) and extend it to demonstrate persistence of this phenotype following *in vivo* EEG recording.

**Fig. 3.**
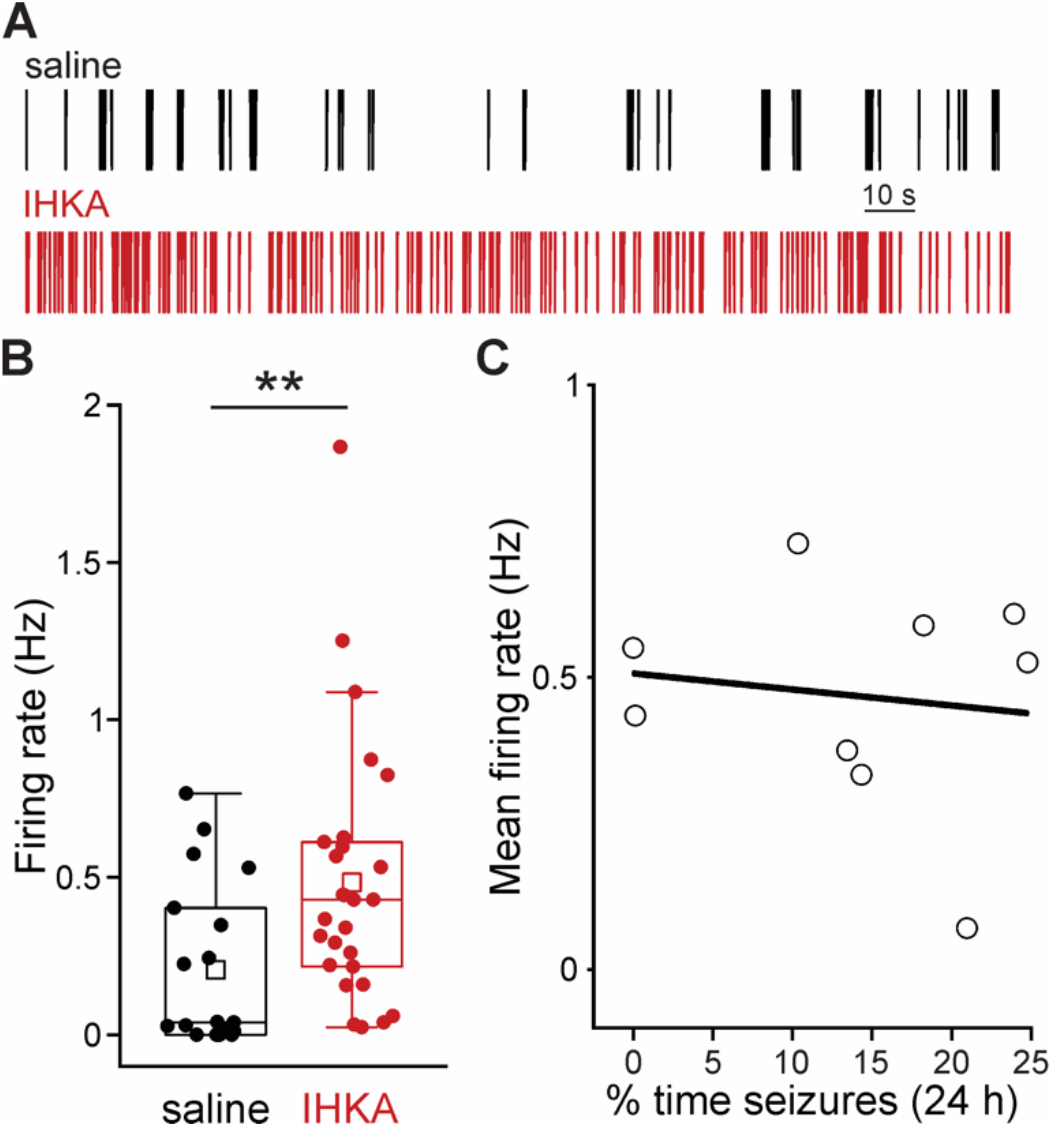
Firing rates of GnRH neurons from IHKA females recorded on diestrus are increased compared to controls following *in vivo* EEG recording through 2 months after injection, but do not correlate with *in vivo* seizure burden. A) Raster plots of example firing activity recorded in loose patch configuration in a neuron from a saline-injected control mouse (top, black) and an IHKA mouse (bottom, red). B) Boxplots for GnRH neuron firing rate recorded in slices from control (black; n=19 cells, 6 mice) and IHKA mice (red; n=27 cells, 9 mice). Boxplots indicate data points within 1.5 times the interquartile range, the mean (squares), the median, and 25–75% range. Circles indicate individual cells. C) Relationship of mean GnRH neuron firing rate in each mouse as a function of the percentage of time in seizure activity in the 24 h prior to slice preparation. Solid line indicates the linear regression of best fit. **p<0.01, Mann-Whitney test

To begin to evaluate the association between chronic seizure burden *in vivo* and GnRH neuron firing rates recorded *in vitro*, the average firing rate was used to represent the GnRH neuron activity level for each mouse and assessed in relation to the percentage of time in seizures in the last 24 hours of EEG recording. In this analysis, no correlations were found between GnRH neuron activity levels and seizure burden (r^2^ = −0.12, p = 0.74, Pearson correlation; **Fig. 3C**).

### 3.3. Increased GnRH mean firing rate is associated with increased variance in individual neuron firing rates

Although mean firing rate provides a snapshot of overall GnRH neuron firing activity in an individual mouse, GnRH neurons have heterogeneous properties (Chu et al., 2010; Constantin et al., 2013; Leupen et al., 2003; Sim et al., 2001; Zhang et al., 2007). Therefore, we evaluated the relationship between the mean firing rate value for each mouse and within-mouse variability in individual cell firing rates in the present data, and found that mean firing rate and standard deviation values for each IHKA mouse were positively correlated (r = 0.67, p = 0.049, Spearman correlation; **Fig. 4A**). This analysis thus supports and extends the previous finding of increased variability of firing rates in GnRH neurons from IHKA females (Li et al., 2018) and indicates that increased mean GnRH neuron activity level in each mouse is associated with increased variance between individual cells, suggesting that subsets of GnRH neurons are particularly susceptible to epilepsy-associated increases in firing activity.

**Fig. 4.**
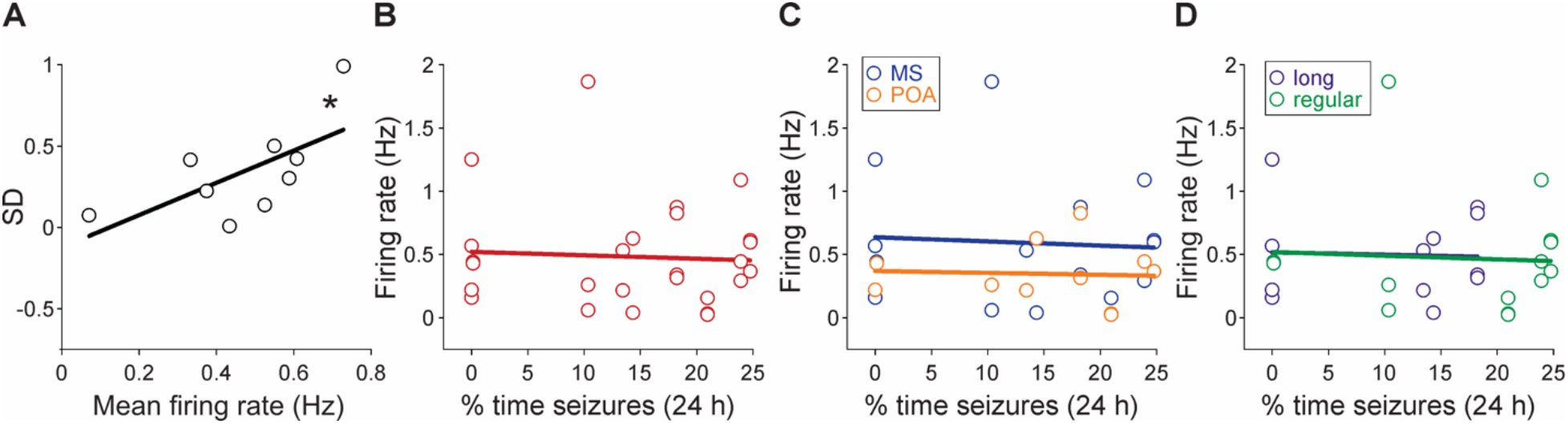
Individual GnRH neuron firing rates are not correlated with *in vivo* seizure burden. A) Variance in GnRH neuron firing rates increases with mean values in IHKA females. Relationship of within-animal standard deviation in GnRH neuron firing rate as a function of the mean GnRH neuron activity level for each mouse. B-D) Relationship of firing rate as a function of the percentage of time in seizure activity in the 24 h prior to slice preparation with all neurons recorded combined (B), separated by soma location in MS (blue) or POA (orange) (C), and separated by classification of estrous cycle as long (≥7-day period, purple) or regular (4-to 6-day period, green) (D). Solid lines indicate the linear regressions of best fit. *p<0.05, Pearson correlations

### 3.4 Lack of correlation between proximal seizure burden and individual GnRH neuron firing rates

Because increased mean firing rate was associated with increased variability of GnRH neuron firing rate values within individual mice, we also performed the correlation analyses evaluating the relationship of *in vivo* seizure burden to GnRH neuron firing activity using individual neuron values rather than mean values for each mouse. These analyses also demonstrated a lack of correlation between these parameters (r^2^ = −0.036, p = 0.76, Pearson correlation; **Fig. 4B**), underscoring that GnRH neuron firing rate was not shaped by *in vivo* seizure burden. Furthermore, we analyzed whether these relationships were influenced by anatomical location of GnRH neuron somata by performing the same analyses with data separated for MS and POA GnRH neurons. However, we did not find correlated relationships of firing activity levels to seizure burden in either subpopulation of GnRH neurons (MS, r^2^ = −0.07, p = 0.82; POA, r^2^ = - 0.096, p = 0.85, Pearson correlations; **Fig. 4C**). Finally, we assessed whether the relationship between MS and POA GnRH neuron firing rates and seizure burden differed depending on estrous cycle length. Neither mice with long (≥7-day cycle period, n=12 cells from 4 mice) nor regular (4-to 6-day period, n=15 cells from 5 mice) cycles, however, showed significant correlations in these parameters (long r^2^ = −0.098, regular r^2^ = −0.07, p>0.8 both groups, Pearson correlations) (**Fig. 4D**).

### 3.5. GnRH neuron firing rates are not different in IHKA females when measured one month after intrahippocampal injection

Spontaneous recurrent seizures have been observed as early as 1-2 weeks after IHKA injection in mice (Bouilleret et al., 1999; Gröticke et al., 2008; Lee et al., 2012; Riban et al., 2002), and IHKA females may not present a seizure-free latent period after injection (Twele et al., 2016). We thus hypothesized that GnRH neuron function may become disrupted much earlier than 2 months after IHKA injection. We previously found, however, that EEG tethering and recording at 1 month after injection produced disruption of the estrous cycle in both controls and IHKA mice (Li et al., 2020), which would complicate interpretation of epilepsy-associated changes in GnRH neuron firing activity. Therefore, we recorded GnRH neuron firing on diestrus at 1 month after IHKA (n = 6 mice) or saline injection (n = 4 mice) without *in vivo* EEG recording prior to brain slice preparation. In contrast to the elevated activity of MS and POA GnRH neurons observed at 2 months after injection (Li et al., 2018) (**Fig. 3**), there was no difference in firing frequency in neurons from IHKA mice (n = 16 cells) compared to controls (n = 9 cells; p = 0.6, Mann-Whitney test) (**Fig. 5**). These findings indicate that although epileptogenesis is typically complete soon after IHKA injection, disrupted GnRH neuron firing in IHKA mice requires longer to develop and is not yet present at 1 month after injection.

**Fig. 5.**
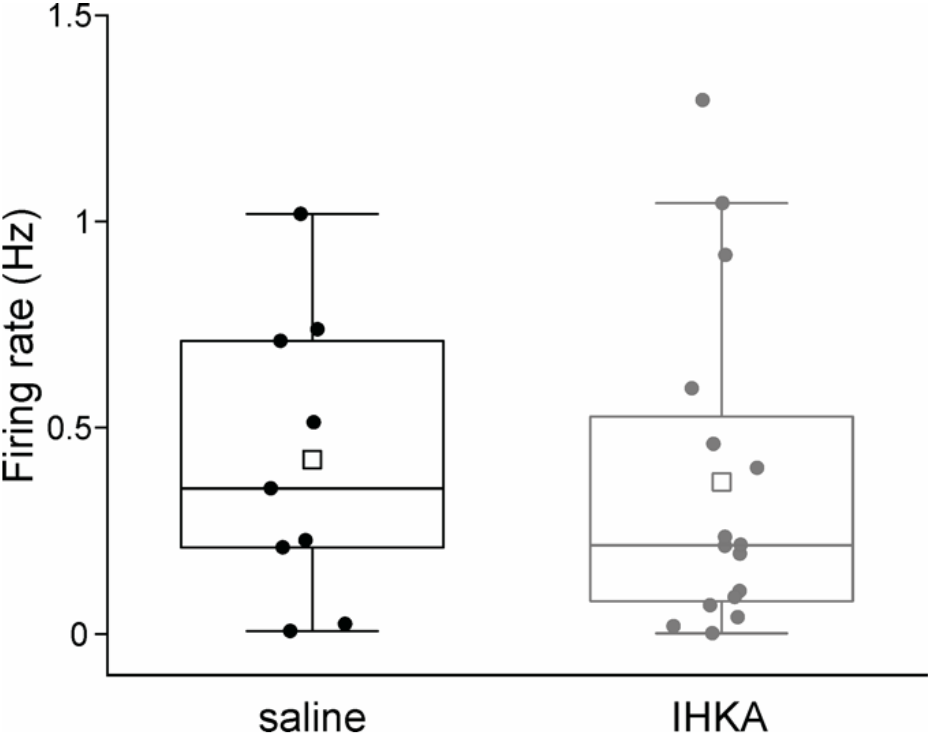
Firing rates of GnRH neurons from IHKA females are not different from controls at 1 month after intrahippocampal injection. Boxplots for GnRH neuron firing rate recorded in slices from control (n=9 cells, 4 mice) and IHKA mice (n=16 cells, 6 mice). Boxplots indicate data points within 1.5 times the interquartile range, the mean (squares), the median, and 25–75% range.

## 4. Discussion

Reproductive endocrine comorbidities are common presentations in people with TLE, but the specific mechanisms that link epilepsy to these comorbidities remain to be elucidated. Rodent models of TLE, in which estrous cycle disruptions in females are common sequelae of seizures and epilepsy in the absence of antiseizure drug treatment, provide useful models for mechanistic investigations in this domain. Moreover, the IHKA mouse model, in which many (but not all) females display estrous cycle disruption, provides a platform to compare the effects of seizures and epilepsy in the presence and absence of such comorbidities. The factors that determine the phenotype of estrous cycle periodicity in IHKA mice remain unclear, but in previous studies we found that increased GnRH neuron firing activity recorded during diestrus was only seen in IHKA mice with long cycles (Li et al., 2018). One possibility is that IHKA females with more severe epilepsy are at higher risk of developing increased GnRH neuron firing activity on diestrus, but this relationship could not be assessed in the prior study as those mice were not recorded in EEG. The present results, however, indicate that the severity of epilepsy and the degree of downstream disruption to GnRH neuron function are not directly correlated.

The present analyses focused on the stage of diestrus as our previous findings indicated a stronger heterogeneity of GnRH neuron firing rates recorded in slices from diestrous IHKA mice compared to those from estrous mice, and that the GnRH neuron firing rates recorded during diestrus, but not estrus, correlated with estrous cycle length (Li et al., 2018). We thus postulated that if GnRH neuron activity levels recorded *in vitro* were correlated to seizure burden recorded *in vivo*, this relationship would be reflected in data collected from diestrous mice. Given the lack of relationship between these parameters as recorded on diestrus, however, in conjunction with the reduced variance of GnRH neuron firing rates recorded on estrus in the previous work, it appears unlikely that such a relationship would be detected in recordings from estrous mice. It should be noted, however, that another time of prominent changes in GnRH neuron firing activity is late proestrus, when a large surge of GnRH release drives a robust surge in luteinizing hormone (LH) release from the pituitary, triggering ovulation (Christian and Moenter, 2010; Czieselsky et al., 2016; Sarkar et al., 1976). The activity levels of GnRH neurons from proestrous IHKA mice have yet to be determined, and it is possible that different relationships between seizure burden and GnRH neuron activity during proestrus may occur, particularly with respect to the timing of seizure activity in relation to the diurnal signals that serve to time the GnRH-LH surge and ovulation to specific times of day in rodents (Bronson and Vom Saal, 1979; Christian et al., 2005; Christian and Moenter, 2010; Everett and Sawyer, 1950; Legan and Karsch, 1975).

Increased firing of GnRH neurons from IHKA males was also observed previously, but with a milder phenotype than that observed in females and effects limited to GnRH neurons with somata in the MS (Li et al., 2018). Moreover, the firing rates of MS GnRH neurons from IHKA mice did not exhibit a sex difference. Whether there is a sex difference in chronic seizure burden in the IHKA mouse model of TLE remains unclear. One study noted increased spontaneous focal hippocampal paroxysmal discharges in male IHKA mice compared to females (Twele et al., 2016), but another study found no sex differences in chronic seizure parameters in IHKA mice (Zeidler et al., 2018). Taken together, the information currently available would indicate that MS-residing GnRH neurons in both males and females may be particularly susceptible to epilepsy-induced changes in firing activity, perhaps reflecting the strong interconnectivity between the medial septum and hippocampus (Gaykema et al., 1991; Mattis et al., 2014; Müller and Remy, 2018; Tóth et al., 1993). This effect, however, appears to be independent of the severity of ongoing seizure burden, and may instead reflect differential intrinsic and/or connectivity properties of the GnRH neurons in this region.

A limitation of the present study is that seizures were only recorded ipsilateral to the site of KA injection in the right dorsal hippocampus. Although seizures in the IHKA model primarily arise in the ipsilateral hippocampal region and many epileptiform events stay focally confined to the hippocampus (Lévesque and Avoli, 2013; Twele et al., 2017), seizure activity can spread and generalize to other brain regions (Sheybani et al., 2018), in some cases producing convulsive seizures (Bui et al., 2018; Lisgaras and Scharfman, 2022; Zeidler et al., 2018). However, other rodent models of TLE with primarily focal seizures also show prominent estrous cycle disruption (Amado et al., 1987; Christian et al., 2020; Edwards et al., 1999; Fawley et al., 2012; Scharfman et al., 2008). Therefore, seizure generalization does not appear to be an absolute requirement for the development of comorbid reproductive endocrine dysfunction, but it remains to be determined whether there is a relationship between the rate of seizure generalization and downstream GnRH neuron and HPG axis dysfunction. Nevertheless, as both electrographic and convulsive seizure activity are consistently detected in the injected hippocampus in IHKA mice (Bouilleret et al., 1999; Lisgaras and Scharfman, 2022; Riban et al., 2002; Sheybani et al., 2018), depth EEG recording in the ipsilateral hippocampus as performed here enables recording of the vast majority of seizures in this model. Therefore, it is unlikely that the present results represent an underestimate of overall seizure burden in these mice.

Clinical evidence linking focal epilepsy to reproductive endocrine disorders indicates that the risk of developing different reproductive endocrine comorbidities in women with TLE depends on the lateralization of seizure focus. Specifically, women with left-sided seizure foci exhibit higher rates of polycystic ovary syndrome, whereas women with right-sided foci are at higher risk of hypothalamic amenorrhea and hypogonadotropic hypogonadism (Herzog, 1993; Herzog et al., 1986; Kalinin and Zheleznova, 2007). In addition, men with TLE show lower circulating testosterone levels than men with seizure foci outside the temporal lobe (Bauer et al., 2004). It is thus possible that seizure focus location, rather than epilepsy severity, is more determinative in shaping the type and degree of reproductive endocrine comorbidities in patients with TLE. In recent studies examining the effect of targeting left or right hippocampus for KA injection in mice, however, we observed similar increases in estrous cycle length regardless of injection site lateralization (Cutia et al., 2021). Likewise, no differences in estrous cycle disruption dependent on left-vs. right-sided seizure focus location were observed in a rat amygdala kindling model (Hum et al., 2009). Therefore, although the relationship of seizure burden and GnRH neuron activity in left-injected IHKA females has yet to be tested directly, it appears unlikely that different results would be found in IHKA mice with left-targeted injections.

The present results also show that GnRH neuron activity is not different in IHKA and control diestrous mice when measured at 1 month after injection, in contrast to the previous and present findings of increased activity in neurons from IHKA mice when recorded at 2 months after injection (Li et al., 2018) (**Fig. 3**). These findings suggest a time course of development of disrupted GnRH neuron function that is distinct from that of epileptogenesis in this model. Furthermore, there may be a potential window of time in which it would theoretically be possible to intervene, even after epileptogenesis is complete, and prevent subsequent changes in GnRH neuron activity. However, the lack of relationship between seizure burden and GnRH neuron firing rates demonstrated in the present studies suggests that a successful intervention of this type would likely either necessitate full cessation of all seizure activity or require a different strategy than seizure suppression alone.

Strong correlations between seizure burden severity and the degree of change in GnRH neuron activity would provide clear links between seizure activity *per se* and dysregulation of hypothalamic circuits controlling reproduction. The present results, however, instead suggest that the propensity to develop reproductive endocrine comorbidities is not merely a consequence of epilepsy severity but instead likely reflects a complex combination of multiple risk factors, such as differences in anatomical connectivity between limbic and hypothalamic structures, intra-hypothalamic circuit function, sensitivity to gonadal steroid hormone feedback, and other possible triggers that are differentially engaged in the presence of seizure activity in certain comorbidity-prone individuals. Although determining these specific mechanisms is beyond the scope of the present work, these analyses provide valuable insight to guide formation of mechanistic hypotheses and future experimental design in this regard.

## Abbreviations

ACSF: artificial cerebrospinal fluid
EEG: electroencephalography
GnRH: gonadotropin-releasing hormone
HPG: hypothalamic-pituitary-gonadal
IHKA: intrahippocampal kainic acid
KA: kainic acid
LH: luteinizing hormone
TLE: temporal lobe epilepsy

## Acknowledgments

We thank Leanna Leverton for assistance with injection surgeries and EEG recordings.

